# The Role of the Basal Ganglia in the Human Cognitive Architecture: A Dynamic Causal Modeling Comparison Across Tasks and Individuals

**DOI:** 10.1101/2025.08.22.671730

**Authors:** Catherine Sibert, Holly Sue Hake, Andrea Stocco

## Abstract

Researchers agree input from the basal ganglia (BG) to the prefrontal cortex (PFC) plays an important role in cognition, but they disagree on its computational properties. Theoretical models characterize the majority of BG input as either direct (directly transmitting information to the PFC), or modulatory (indirectly influencing PFC activity through the gating of signals from other cortical areas). To determine the computational nature of these BG-PFC inputs in cognition, we tested three alternative connectivity configurations (Direct, Modulatory, and Mixed) within a large-scale cognitive architecture. This architecture, the Common Model of Cognition, has been independently validated using fMRI data from the Human Connectome Project (HCP). The Direct model reflected the standard CMC configuration, featuring a bidirectional, direct connection between the BG and PFC modules. The Modulatory model removed this direct link, instead incorporating a unidirectional connection from the PFC to the BG and two modulatory pathways from the BG to the PFC that passed through other cortical modules. The Mixed model included both direct and modulatory connections. Using fMRI data from 200 HCP participants performing six cognitive tasks and one resting-state session, we applied Dynamic Causal Modeling (DCM) to estimate and compare the influence of these different BG–PFC connectivity patterns. Here, we show that for each of the six cognitive tasks and resting state, the Mixed model consistently outperformed the Direct and Modulatory models. The next best model depended on the specific cognitive task, suggesting the ability for the BG to flexibly adapt to various task demands. Taken together, the current data provide evidence for a likely set of core computations that the BG uses to differentially regulate cortical activity.

**Author Summary:** This study aimed to understand the nature of regulatory input from the basal ganglia to the prefrontal cortex in cognition using a neuro-computational approach. Three model architectures of basal ganglia connectivity were tested using fMRI data from 200 individuals during cognitive tasks and resting state sessions. Our findings indicate that the Mixed architecture, in which the basal ganglia can affect the prefrontal cortex both directly and indirectly, outperformed the Direct and Modulatory architectures across various cognitive tasks and resting state sessions. The results suggest that the basal ganglia likely uses a set of core computations to regulate cortical activity, which is present during task and task-free behavior.

## Introduction

“My basal ganglia’s an A to the K” –Eminem, *Greatest*, 2018

The basal ganglia (BG) are a set of interconnected subcortical nuclei surrounding the thalamus that receive inputs from nearly the entire cortex [1], and project to the prefrontal cortex through the dorsal and medial nuclei of the thalamus. Historically, the BG have been closely associated with motor functions, especially in neurodegenerative pathologies such as Parkinson’s disease and Huntington’s diseases. Even in these cases, however, non-motor symptoms can be widespread and prevalent, and can include a decline in working memory, executive function, and attention [2]. Furthermore, abnormal BG function has been implied in a variety of other pathologies, including obsessive-compulsive disorder [3], Tourette’s syndrome [4], and attention deficit disorder (Shaw et al., 2014). The scope of BG involvement in cognition has further increased with fMRI studies: at the time of writing this paper, more than 4,000 BG papers can be found in Neurosynth (neurosynth.org), spanning a variety of functions including attention [5], language [6], learning, working memory [7], and memory retrieval [8].

In summary, neurological and neurocognitive research have demonstrated a widespread involvement of the BG across multiple core cognitive functions and mental disorders. This variety suggests two things. First, understanding the function of the BG is essential for correctly understanding a variety of neuropathologies, and second, the BG likely provides a set of core computations that are used by multiple specialized circuits in the human brain.

### Computational Models of the Basal Ganglia

A number of computational models have been put forward to conceptualize and formalize the core computations of the basal ganglia, growing in complexity and biological fidelity from early actor-critic models [9] to complex and integrated spiking neuron systems [10,11]. Despite the variety of domains that have been tackled by these models and the differences in their approaches, models have converged on a series of shared assumptions. For example, all models agree that inputs from the cortex undergo some form of integration and selection in the striatum. They also agree that dopamine carries a reward signal, usually framed as the reward prediction error in reinforcement learning theory [12,13], although alternative views have been proposed [14,15]. Additionally, all models agree that dopamine drives plasticity and results in synaptic potentiation and learning. Finally, all models agree that the outputs of the BG circuit (through the thalamus) have profound and measurable effects on the neural activity of the prefrontal cortex.

Despite their convergence, models disagree about how this influence is exerted. Specifically, whether this influence is exerted directly, indirectly, or both. In this paper, we examine how BG activity affects PFC responses within the broader context of cortical dynamics. Rather than isolating BG–PFC interactions, we embed them within a large-scale brain architecture to assess how BG output shapes PFC activity in the presence of other ongoing cortical processes.We argue that, in general, these effects can be divided into two categories: direct and modulatory. In the majority of models, the effects of the BG on a cortical region are local, and can be observed and modeled without any reference to the state of other cortical regions. Being local, these effects can be categorized as direct. Examples of these models include models of action selection [16–18], models motor control [19] and some models of working memory [20].

In other models, however, the effect of BG outputs is to control how signals flow *between* cortical areas. In statistical terms, the activity of the BG is best seen as a modulator of the signals travelling between their PFC target and other cortical regions. Therefore, we refer to this effect as modulatory. Models of this type include the PBWM model of working memory [21,22], and the SPAUN neural architecture [10]. In the PBWM, for example, cortical areas maintain information by default, resisting inputs from other dedicated regions. The BG switches the states of PFC cells, allowing them to cease the maintenance of the current pattern of activation, and instead receive inputs from a posterior cortical region. In other words, the BG performs a “gating” operation, controlling the flux of inputs from posterior to prefrontal cortical regions.

The distinction between “direct” and “modulatory” models can be blurred, and some models include both types of effect. For example, in the Conditional Routing model [23], the BG is initially used to resolve conflicts and route information from posterior regions to the PFC; however, with learning, the BG can learn to directly affect PFC activity without the need for a posterior input.

### Dynamic Causal Modeling

To test whether the effects of the BG are direct, modulatory, or both, one needs to examine the *effective* connectivity between the BG and other brain regions; that is, the extent to which changes in one region affect other regions. One elegant way to do this is through Dynamic Causal Modeling (DCM; [24]). This framework captures effective connectivity as a set of parameters that estimate the extent to which the timeseries of one brain region affects the rate of change of another. The change in the output activity **y** of *N* regions is captured here as a differential equation:

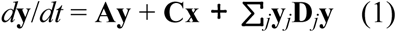

In this equation, **x** is a binary vector that defines the presence of each of *M* task conditions at any moment in time and **y** is a vector containing the current activity of all *N* regions. Effective connectivity is represented by four parameter matrices: **A** is an *N*-by-*N* matrix of parameters that capture the intrinsic connectivity between regions, **C** is an *M*-by-*N* matrix that defines the effects the *M* task conditions on each region, and **D** is an *N*-by-*N-*by-*N* matrix that defines the modulatory effects that regions have on the connections between other regions [25].

The most critical feature of the DCM framework is its ability to model both first-order and second-order interactions. First-order interactions refer to direct influences between regions—how activity in one node affects another—captured by matrix **A**. Second-order interactions represent modulatory effects, where a region influences the strength of connectivity between two other regions, as captured by matrix **D**. This is of particular interest as it allows for the capture of the hypothesized nature of BG functionality, whether *direct* or *modulatory*.

### Underlying Brain Architecture

One of the downsides of DCM is that, instead of providing data-driven estimates of network structure, it can only measure connectivity within a given network, defined by the matrix **A**. In turn, this means that the precise measure of connectivity between two regions might change depending on the larger network context in which they are embedded. Because we are carrying out an investigation of the general role of the basal ganglia across multiple tasks, it is essential to apply DCM within a reduced but biologically-plausible network that captures the essential characteristics of the human brain’s architecture. Such a network architecture would need to be large enough to span the most common regions, yet simple enough to make DCM computationally tractable.

In this paper, we have chosen to rely on a specific architecture that had been previously tested in DCM applications and shown to possess the desired qualities. The architecture is based on the Common Model of Cognition (CMC: [26]), a conceptual framework designed to distill the lessons learned from 40 years of cognitive science research on the essential modules of the mind and their relationships. Abstract computations are categorized into five functional components and the connections between them are specified in a graph structure (Fig 1).

**Fig 1.**
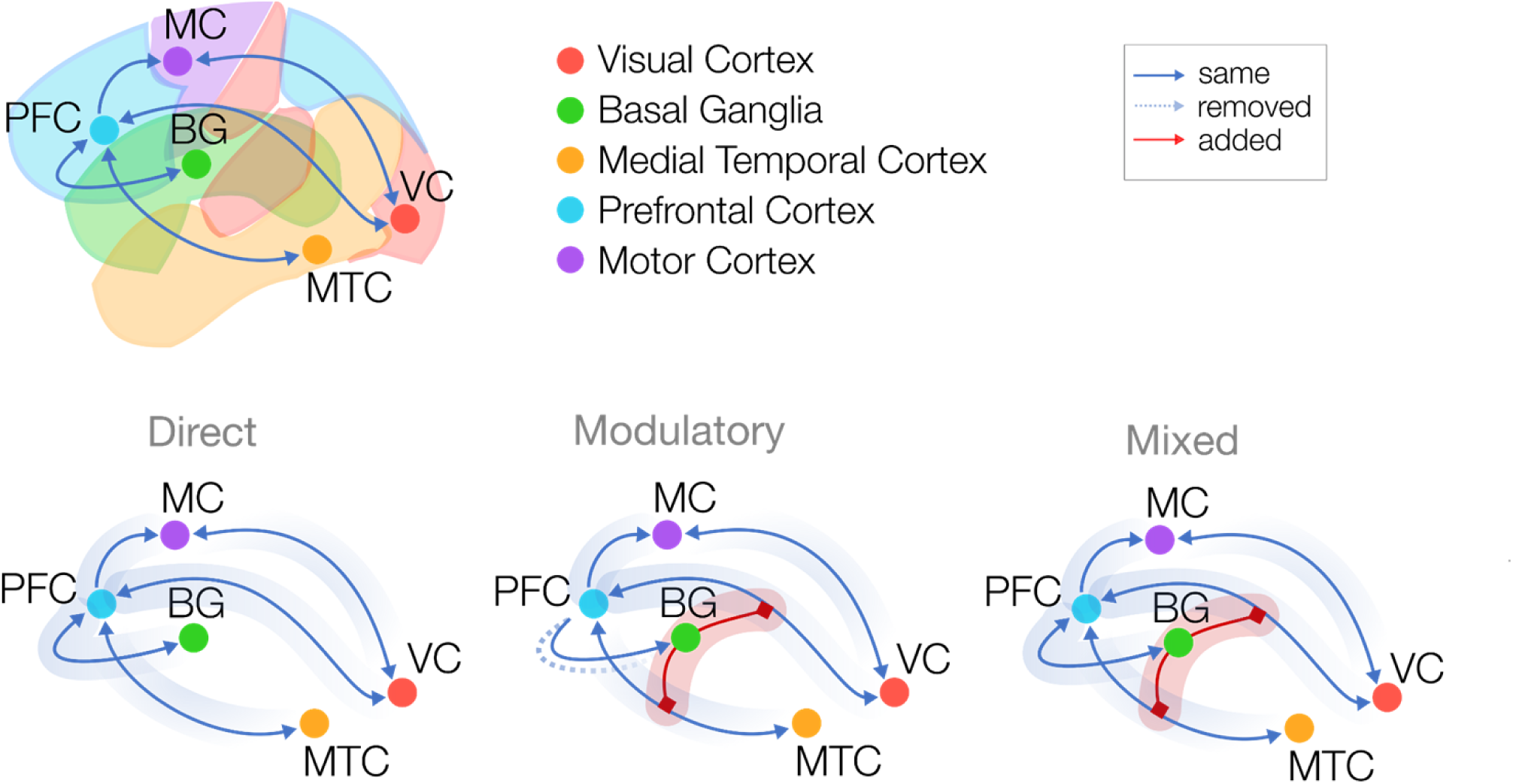
Models capturing three different views of basal ganglia function. The structure of all three network architectures is illustrated. The top panel shows the anatomical layout, with key regions labeled: Visual Cortex (VC), Basal Ganglia (BG), Medial Temporal Cortex (MTC), Prefrontal Cortex (PFC), and Motor Cortex (MC). The bottom panel presents the Direct (left), Modulatory (center), and Mixed (right) network models, depicting information flow pathways and their connectivity structures. Solid lines represent direct excitatory pathways, dashed lines represent connections that were removed, and red lines represent modulatory connections.

Although it was not proposed specifically as a brain architecture, the underlying assumptions of the CMC are shared by the most popular architectures used to model behavioral and neural data in cognitive tasks [10,27,28]. Furthermore, a number of studies have found that the CMC is surprisingly effective at modeling brain activity across tasks and individuals [29–31]. In this interpretation, the CMC’s five functional components (long-term memory, working memory, procedural memory, perception systems, and action systems) are mapped onto large-scale brain regions (medial temporal lobe, prefrontal cortex, basal ganglia, primary and visual cortex, and motor cortex) and their relations are translated into predicted patterns of functional connectivity, with the PFC playing the role of a central hub while allowing direct connections from visual to motor areas. In other words, the neural counterparts of the functional components and their connections serve as a simplified architecture for the human brain. Importantly, the neural interpretation of the CMC explicitly includes the BG as one of its components and serves as the neural implementation of the “procedural memory” component.

Recent work has shown that this architecture provides a remarkably good fit to fMRI data from over 200 participants across a variety of representative tasks [29]. Converging evidence for the validity of the CMC as a realistic high-level model of the brain’s architecture also comes from other two sources. First, the same architecture can be shown to hold not only when analyzing task-based activity, but also when analyzing resting state data [32]. Second, the same architecture emerges when the original time-series data from each region is analyzed using Granger causality [33]. Taken together, these findings confirm that the CMC could be used as a high-level approximation of the brain’s functional connectivity design.

Inspired by its success in accounting for functional connectivity, we decided to extend this approach to investigate the role of the BG by creating three different variants of the CMC architecture, corresponding to three different characterizations of the BG. In the Direct model, the BG directly affects the PFC (corresponding to the “working memory” component of the CMC). In the Modulatory model, the BG does not affect the PFC directly, but instead mediates the connectivity between perceptual inputs and long-term memory to the PFC. Finally, a third, Mixed model was examined; in this model, the BG can exert both direct and modulatory effects on PFC. Note that, because the Mixed model contains all of the connections and the parameters of the other two, the Direct and Modulatory models can be said to be nested under the Mixed model.

### The Human Connectome Project Dataset

Previous studies have examined direct and modulatory BG connections, but only across a narrow range of tasks [30,31]. Because these models embody different hypotheses about the general functional role of the BG in cognition, it is important to test them across multiple tasks. This is particularly important when considering the Mixed model, as the influence of direct and modulatory connections might adaptively change depending on the situation, and testing a single task might yield biased results that are not representative of the whole functionality. Although previous attempts have found that the Modulatory model fits the data better than the Direct model within a specific task [30] and in healthy participants over BG-impaired patients [34,35], a true test of the models requires an examination of a larger variety of tasks that span different cognitive domains.

To this end, we used a subset of the fMRI data available from the Human Connectome Project [36], one of the largest high-quality human neuroimaging datasets. The HCP includes data from 1,200 young adults performing different tasks, six of which were selected to cover different domains of cognition. Table 1 lists these six paradigms and their relevant experimental conditions. The three architectural models described above were translated into equivalent DCM network models, and data from the six tasks were used to fit network connections that best predicted the observed human brain activity. These predictions were compared across the alternate model structures.

**Table 1.** Paradigms used in the HCP dataset.

| Paradigm | Condition (Baseline) |
| --- | --- |
| Emotion Processing | Fearful and angry faces (Neutral faces) |
| Incentive Processing | Winning blocks of choices (Losing blocks) |
| Language and Math | Answering questions about stories or math (Passive listening) |
| Relational Reasoning | Inferring the rule in relational arrays (No rule in control arrays) |
| Social Cognition | Socially interacting shapes (randomly moving shapes) |
| Working Memory | 2-Back blocks of stimuli (0-Back) |
| Resting State | 15 min eyes open fixation point |

## Results

### Likelihood-base Model Comparison

Given two or more generative models that produce the same data **x**, it is possible to compare them by estimating a likelihood function *L*(*m* | **x**) and select the model with the highest likelihood. A model’s likelihood is is defined as the probability that *m* would generate **x**, that is, *L*(*m* | **x**) = *P*(**x** | *m*). Group-level likelihood values for a model m can then be expressed as the product of the likelihood of that model fitting each participant *p*, i.e., ∏*_p_ L*(*m* | **x***_p_*) [37]. Because probability values might become vanishingly small (leading to rounding errors), it is customary to compare models in terms of their log-likelihood, with the group-level log-likelihood then becoming the sum of all of the individual log-likelihoods: ∑*_p_* log *L*(*m* | **x***_p_*). Although more sophisticated model comparison procedures have been proposed [37], the log-likelihood-based metric used here is not only the most easily interpretable but also the most relevant, as it specifically applies to cases in which it is assumed that the model is constant or architectural across individuals [39].

Figure 2 illustrates the group-level log-likelihoods of the three models in each of the six tasks. This represents the likelihood that a particular model will provide the best fit for a random subject within the sample. Higher log-likelihoods correspond to a higher probability that a model is capturing the actual dynamics of cognition. Because the group-level log-likelihoods vary dramatically across tasks and models, the figure presents them as *relative* log-likelihoods: within each task, the lowest log-likelihood value is subtracted from all the others. As a result, the worst-fitting model always has a relative log-likelihood value of zero, while the other two are positive. In all tasks, the Mixed model significantly outperforms the other two.

**Fig 2.**
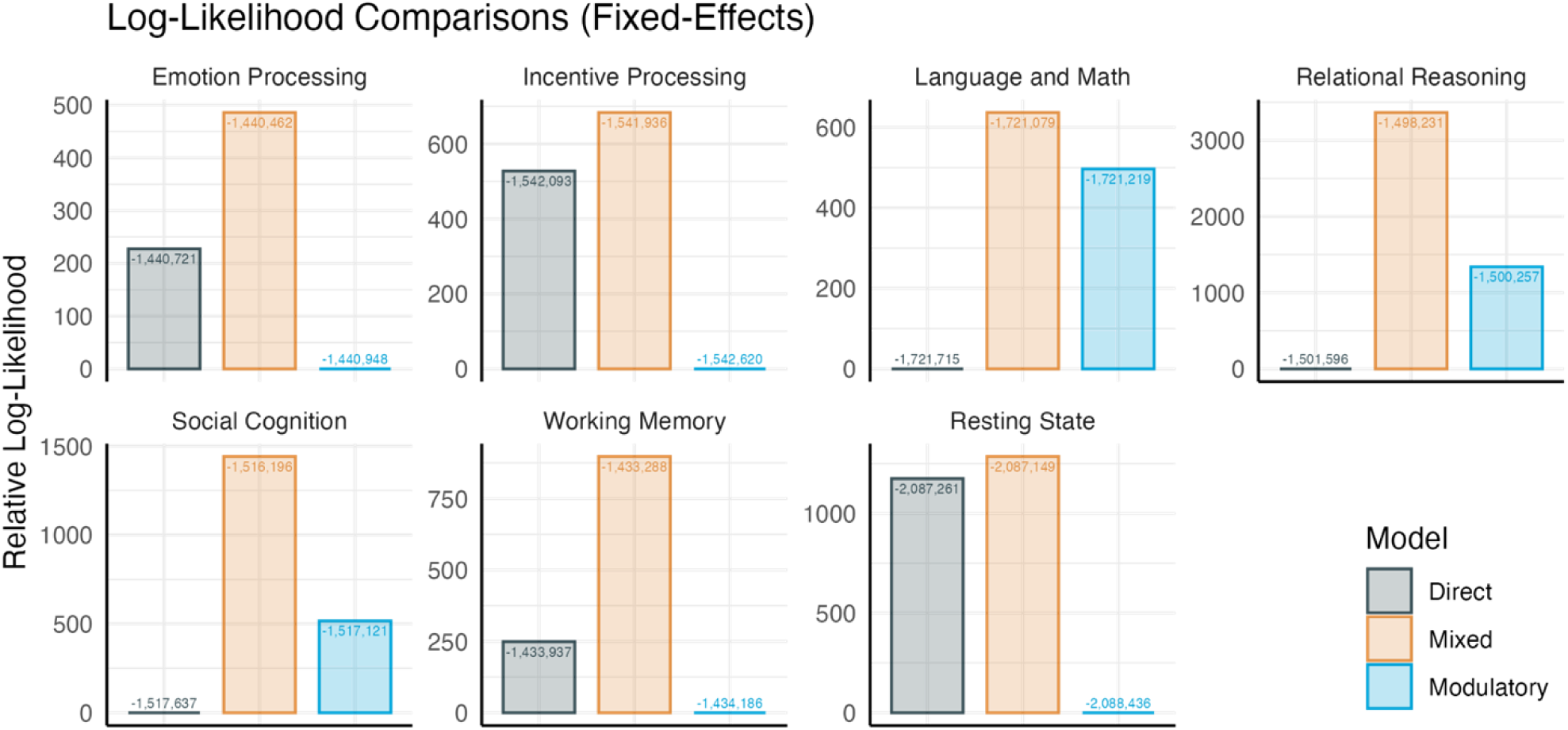
Relative log-likelihoods across tasks. Bars represent the relative log-likelihoods of each model for each task; numbers represent the absolute likelihood values.

The results also show significant differences across tasks. In the Relational Reasoning task, for example, the Mixed model dramatically outperforms the other two. This is to be expected, given that this task requires the integration of knowledge. It is also interesting to note the relative fit of the other two models is dependent on the task: while the Direct model is preferred over the Modulatory model in the incentive and emotion processing tasks (both of which require the integration of reward signals), the Modulatory outperforms the Direct model in the Relational and the Language and Math paradigms.

### Accounting for Differences in Complexity

Since the three models differ also in terms of the number of parameters, it is possible that the Mixed model’s greater likelihood is due to it simply having more degrees of freedom to fit the data. Although common corrections can be applied (such as BIC and AIC) for the number of parameters, the fact that the Direct and Modulatory models are nested within the Mixed model allows us to use Wilks’ theorem [40], which accounts for the different number of parameters and translates the log-likelihood difference into interpretable *p*-values. The theorem states that, for two models of which one is nested, the probability that the fit of the more complex model is due to chance (its *p*-value) approximates the probability of obtaining the value of 2log(λ) (twice the log-likelihood difference) in a χ^2^ distribution with degrees of freedom corresponding to the difference in the number of parameters. Using this theorem, we calculated the probability that the greater fit of the Mixed model is due to chance (note that this comparison accounts for the greater complexity of the Mixed model in the χ^2^ distribution). Figure 3 illustrates the performance of the Mixed model against the Direct model (top) and the Modulatory model (bottom). In the figure, the shaded curves depict the corresponding χ^2^ distributions (note that the distributions are the same for all tasks since they depend only on the differences in parameters between the two models), and the colored points correspond to the value of the 2log(λ) statistic for all the tasks. As the figure shows, all of the differences in log- likelihood are far to the right of χ^2^ distributions, corresponding to *p* < .0001 for all comparisons in all tasks. This implies that the Mixed model’s superiority cannot be accounted for by its greater complexity.

**Fig 3.**
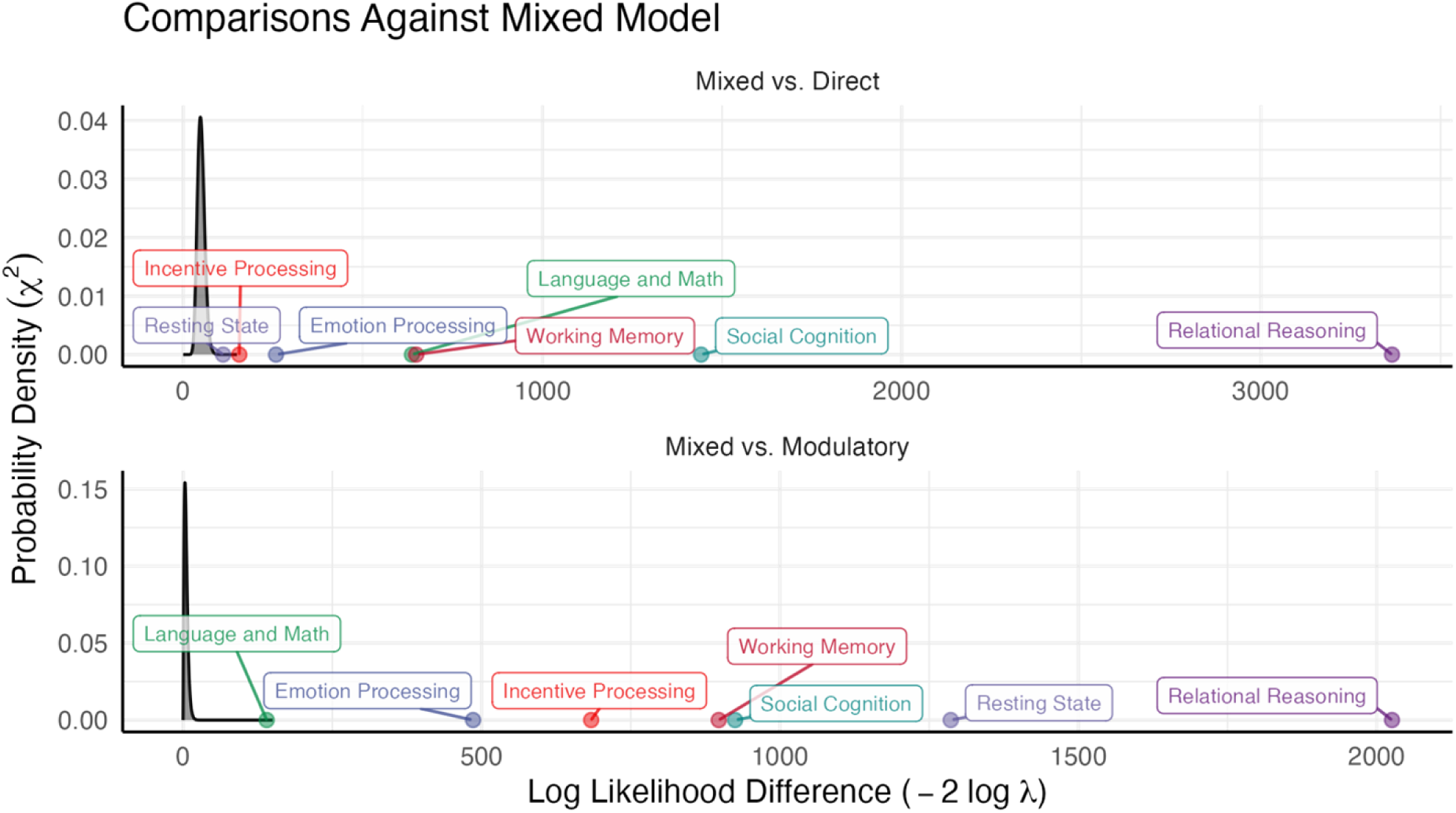
Model comparison, accounting for complexity using Wilks’ theorem. Adjusted likelihood that the Mixed model would outperform the Direct model (top) or the Modulatory model (bottom) across all tasks.

### Accounting for Architecture Variability Across Individuals

Finally, both of the previous analyses assume that the “correct” model is a latent architecture common to all participants, and that differences across participants are in fact measurement noise. Statistically, this assumption is equivalent to assuming that the architecture is a fixed effect. It is possible, however, that different individuals show differences in neuroanatomy that are, in fact, consistent with different basal ganglia architectures; in statistical terms, this is analogous to treating the underlying architecture as a *random effect*. Friston et al [37] provided a method for such a random-effect analysis of DCM models, modeling the probability that each of the three models would best fit a participant is modeled as a Dirichlet distribution. The result of this method is a posterior probability distribution for each model, with the distribution values representing the uncertainty about the probability that a given model would fit a random individual from the population. Figure 4 provides visual representations of these distributions for all three models in each of the six tasks and the resting-state condition. As the figure makes it clear, the distributions for the Mixed model are consistently centered around higher values than the other two models. The relative fit of each model can be quantified by computing their exceedance probabilities, that is, the probability that a value from a model’s distribution would be greater than that drawn from any of the other two models (labels in Figure 4). The exceedance probabilities of the Mixed model ranges between 59.5% and 100%. Thus, even after relaxing the assumption that the models capture an invariant underlying architecture and conducting a random effects analysis, the Mixed model still emerges as the best-fitting model across all tasks.

**Fig 4.**
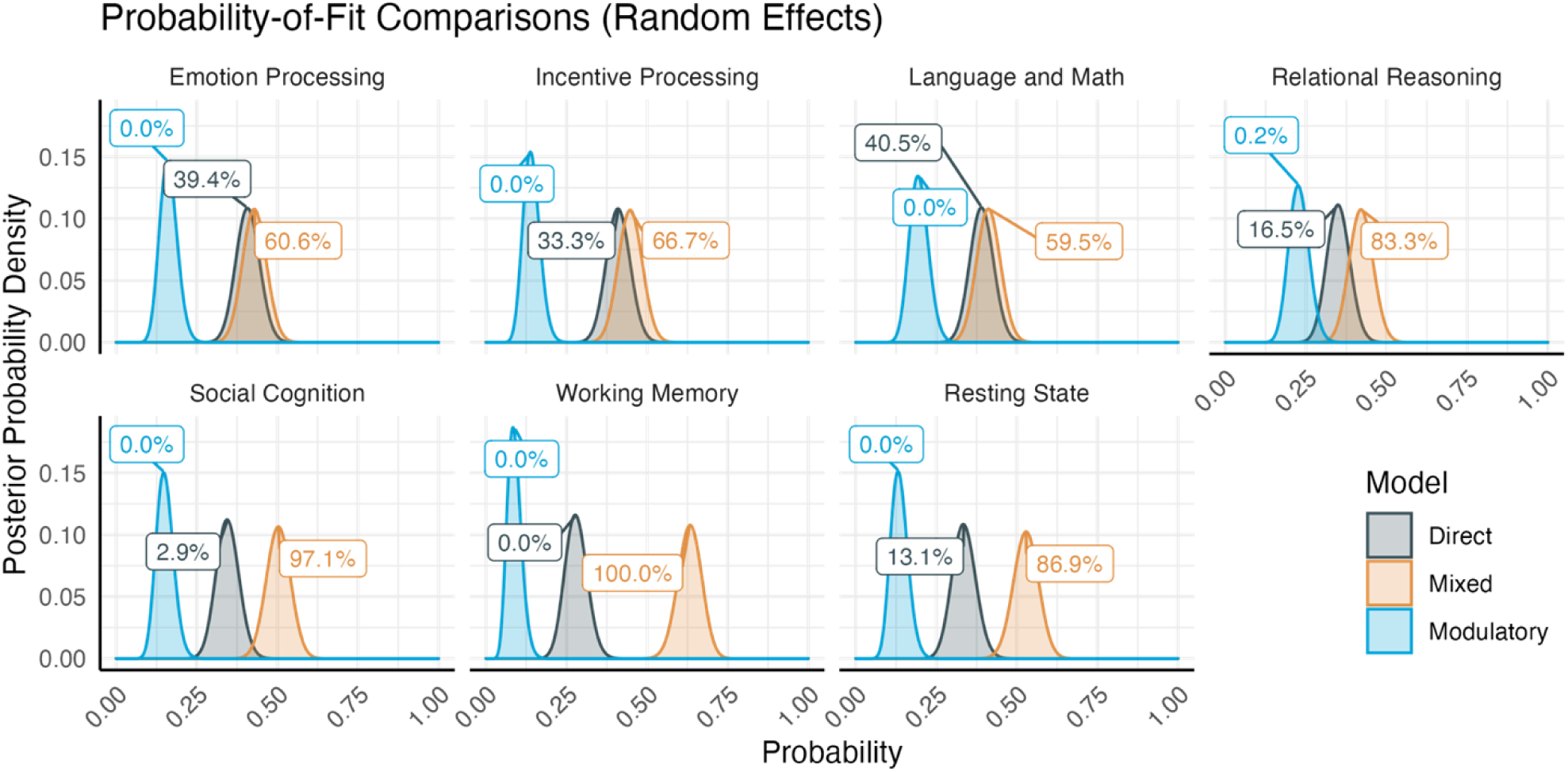
Posterior distributions of the probability that each of the three possible models (Direct, Modulatory, and Mixed) would fit a participant in each of the experimental paradigms. Labels indicate exceedance probability which reflect the relative fit of each model.

### Effective Connectivity Values

Although the previous analyses converge in affirming that a Mixed architecture (including both direct and modulatory effects) best explains the nature of the BG’s input to the PFC, we still need to confirm whether all of the connectivity parameters in the Mixed model are equally important. To do so, we averaged the values of each connection in the Mixed model across all participants for any given task using a Bayesian parameter average method. These connections included the BG to PFC pathway from the Direct model, and the BG to PFC via the Medial Temporal Cortex, and BG to PFC via the Visual Cortex from the Modulatory model. In this procedure, the precision (that is, inverse of the standard deviation) of the final probability distribution is the sum of the precisions of each individual observation, and the mean average is the mean of each observation weighted by its precision. This procedure yields two measures: the expected value of the parameter and the corresponding posterior probability that such value is greater than zero, which are depicted and reported in Figure 5.

**Fig 5.**
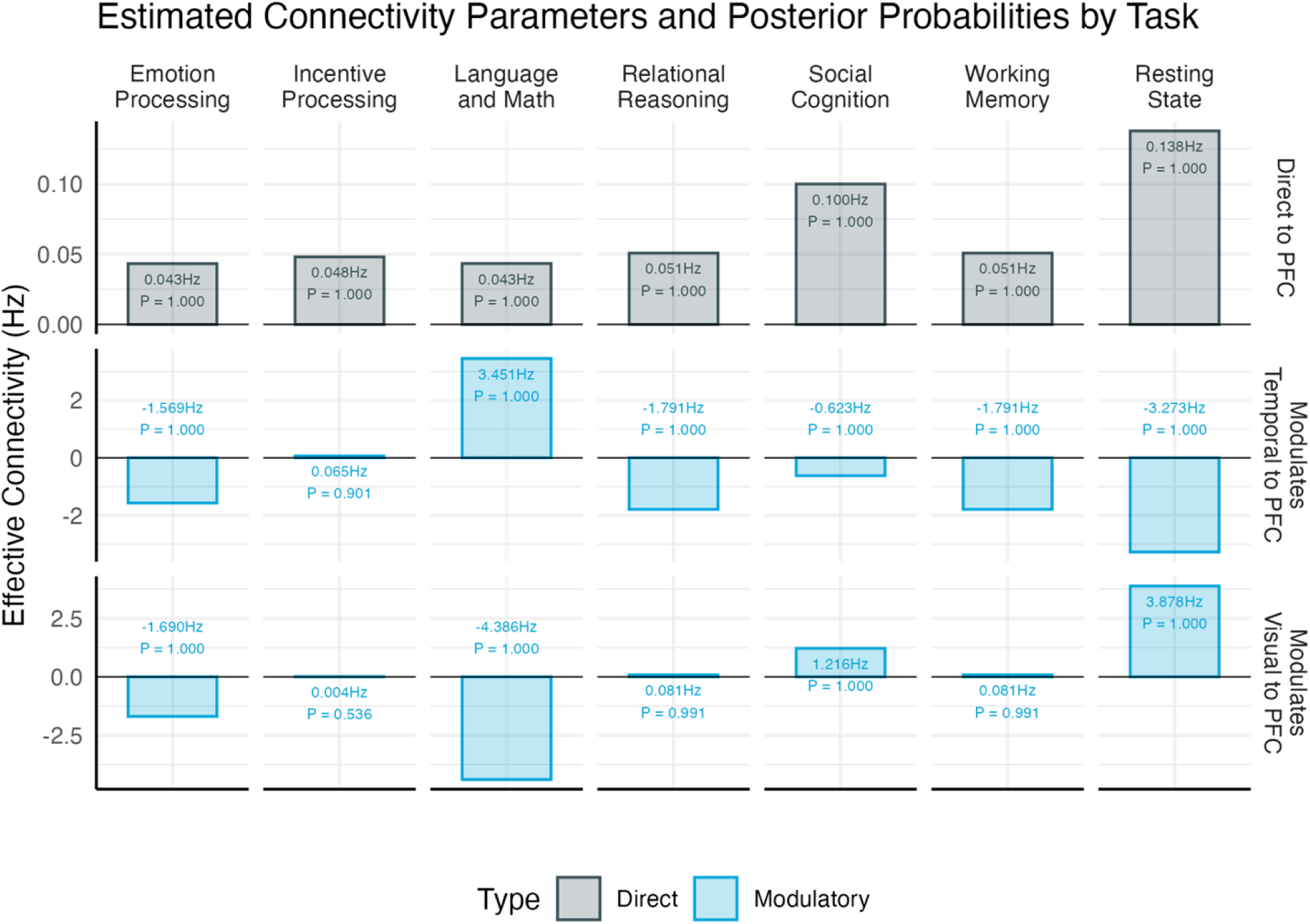
Mean estimated values of the direct (grey) and modulatory (blue) connectivity parameters across all paradigms. The numbers above each bar represent the mean connectivity value and the associated posterior probability that the value is different from zero.

As the figure shows, all three connections are significant across tasks with a posterior probability greater than *P* > 0.9. The only exception is the connection that modulates visual input to PFC, which fails to reach significance in the Incentive Processing task (it is still significant in all other tasks).

## Discussion

The basal ganglia has been linked to a wide variety of cognitive functions, but to date, there has been no consensus of the functional role that it plays within a broader cognitive structure, with particular emphasis on the connection between it and the prefrontal cortex. Existing computational models categorize the effects of the BG as either local and direct, or involving other regions (i.e. indirect or modulatory). Based on the previous success of using Dynamic Causal Modeling to compare competing accounts of cognitive architecture structure, the same method was used here to formulate and compare alternate accounts of BG connectivity. All models were variations of the Common Model of Cognition, an abstract blueprint for cognition that was previously effective at capturing neural activity across multiple participants and tasks. Hypotheses proposing a local and direct connection between the BG and the PFC were represented by the Direct model, which is also the base CMC model. In this structure the BG, represented by the Procedural Memory CMC component, is directly connected to the PFC, represented by the Working Memory CMC component. Hypotheses proposing the involvement of other brain regions were represented by the Modulatory model. In this structure, the direct connection between the BG and the PFC is removed (though the direct connection in the reverse direction from the PFC to the BG remains in place), and activity in the BG instead affects the magnitude of the incoming direct connections from other regions to the PFC, in this case from the Long Term Memory CMC component (Medial Temporal Cortex, MTC) and the Perceptual CMC component (Visual Cortex, VC). A third model, the Mixed model, includes both the direct connection between the BG and the PFC, as well as the modulatory connections modifying inputs from MTC and VC. To test which best represents how neural activity is propagated from the BG to the cortex, a large fMRI dataset of 200 participants performing six, representative cognitive tasks was analyzed through Dynamic Causal Modeling. The comparison showed that a Mixed model that includes both direct and modulatory connectivity consistently outperformed models that include only direct or only modulatory connections. It was also found that the relative rankings of the direct and modulatory models depended on the specific task, suggesting that BG is a flexible system that adapts to task demands.

These results represent a promising use case for the use of comparative models of functional connectivity for the exploration and validation of theoretical notions of brain functionality —although caution is certainly needed when interpreting regional activity within a network [40]. While previous models take an either-or approach to explaining BG connectivity (fully direct or fully modulatory), the results of this model provide evidence that a mixed approach may better represent the true nature of the BG.

The results are silent, however, about the exact nature of the BG computations and the relationship between direct and modulatory connectivity. It is unclear, for instance, if both connection types are being used all the time for every task, or if reliance on a particular connection type depends on a particular task type or environment. The Conditional Routing Model [23] has proposed a useful framework for approaching this uncertainty, in which tasks initially require modulation but, with practice, switch to a more direct effect. Following the completion of the study described above, a very basic attempt to explore this possibility was done by comparing models fit to the data during the first half of a session against models fit to data from the second half of the session. If the nature of BG connectivity changes as a result of learning, we would expect to see significantly different weights for the BG connections during each half of the trial. However, we did not observe a consistent pattern of changes to weight strength in our tested sample. This should not discount the Conditional Model as a possibility; more likely our analysis was not sensitive enough to capture learning effects, or these effects were not sufficiently pronounced as to have an impact within a single 15-minute session of a simple task. Further model-based experiments may be able to better capture the relationship between the BG and learning. Finally, the results show the value of the CMC as a potential, high-level brain architecture.

## Methods

The study presented herein consists of an analysis of neuroimaging data from a subset of *N*=200 individuals in the Human Connectome Project.

### fMRI data

The original HCP task-fMRI dataset includes two resting-state runs and seven task-based runs, each designed to probe a distinct facet of cognition through different experimental paradigms. Our analysis included six of these seven tasks, excluding the motor localization task due to its limited cognitive demands and lack of higher-order cognitive engagement. We also included data from only the first of the two resting-state sessions. A detailed description of all tasks and the rationale for their selection can be found in the original HCP papers [36,41].

### Ethics Statement

Since the study involved the analysis of data that had been previously connected from *N* = 200 participants, the Institutional Review Board at the University of Washington determined that no special approval was needed. The original study’s ethics review and approval can be found in [36].

### Data processing and analysis

#### Imaging acquisition parameters

As reported in Barch et al., [41], functional neuroimages were acquired with a 32-channel head coil on a 3T Siemens Skyra with TR = 720 ms, TE = 33.1 ms, FA = 52°, FOV = 208 × 180 mm. Each image consisted of 72 2.0mm oblique slices with 0-mm gap in-between. Each slice had an in-plane resolution of 2.0 × 2.0 mm. Images were acquired with a multi-band acceleration factor of 8X.

#### Image preprocessing

Images were acquired in the “minimally preprocessed” format [36], which includes unwarping to correct for magnetic field distortion and the removal of physiological noise artifacts. The minimally preprocessed images were then de-spiked and re-aligned to the first image in each run. Additional motion-related artifacts were removed through linear regression, using the affine (*x*, *y*, *z*) and rotation (pitch, yaw, roll) components estimated from the re-alignment step, plus their temporal derivatives. The images were then normalized to the MNI ICBM152 template and smoothed with an isotropic 8.0 mm FWHM Gaussian kernel.

#### Canonical GLM

A canonical GLM analysis was conducted on the preprocessed data using a mass-univariate approach as implemented in the SPM12 software package. First-level (i.e., individual-level) models were created for each participant. The model regressors were obtained by convolving a design matrix with a hemodynamic response function; the design matrix replicated the analysis of Barch et al., (2013), and included regressors for the specific conditions of interest described in Table 1. Second-level (i.e., group-level) models were created using the brain-wise parametric images generated for each participant as input.

#### DCM-specific GLM

A second GLM analysis was carried out to define the event vectors **x** that Eq. 1 provides as a formal representation of external (or, in the case of resting state, intrinsic) drivers of neural activity. Because these models are not used to perform a standalone data analysis, the experimental events and conditions are allowed to be collinear.

All of the HCP paradigms include a critical condition and a baseline (Table 1). As is common in DCM analysis, these two experimental conditions were modeled in a layered, rather than orthogonal fashion, with all trials included in a generic regressor and trials from the critical condition representing a special regressor that further drives neural activity. Following the procedures of Stocco et al., (2021), the association between regressors and ROIs was kept constant across all tasks, with the generic regressor affecting the perceptual component and the critical regressor additionally affecting WM,capturing the greater mental effort that is common to all critical conditions (see Fig. 6).

**Fig 6.**
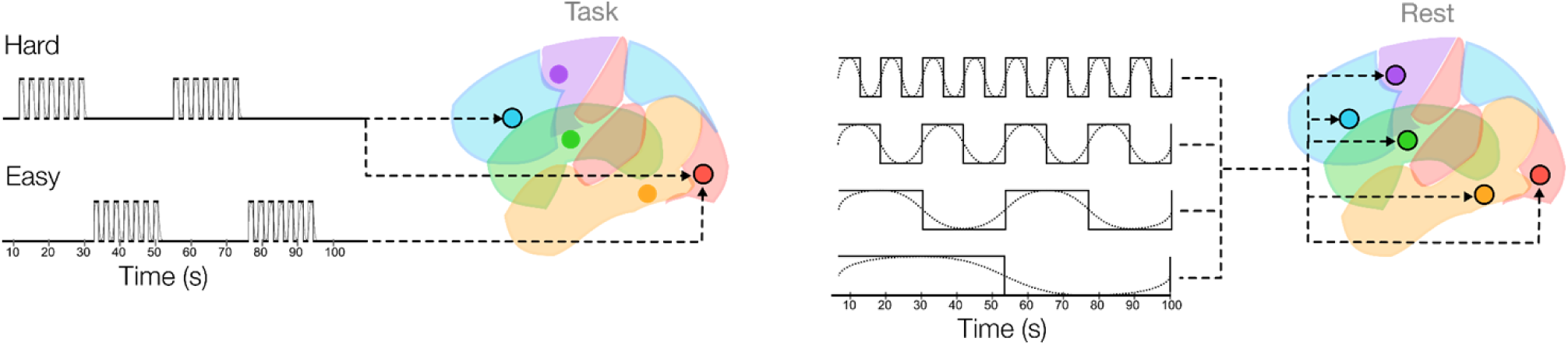
Task and resting-state input designs for model estimation. Left: During task sessions, “Hard”, or critical conditions simultaneously drive perception and working memory systems, while “Easy”, or baseline conditions engage only perceptual regions. Right: During resting-state sessions, “Artificial” inputs are applied uniformly across all modeled regions to enable parameter estimation. Dotted sine waves indicate the timing and duration of input drives.

In the case of resting state data, eight artificial regressors were generated to capture the spontaneous low-frequency oscillations in every node of the network. Following [32,42] these oscillatory regressors were generated by first creating four sine and cosine waves at different representative frequencies (0.01, 0.02, 0.04, and 0.08 Hz, respectively), and then discretizing them to make them suitable for dynamic causal modeling (Fig. 6).

#### Task-based Regions of interest

Regions of Interest (ROIs) for each task and participant were defined using the method described in [29]. For each CMC component, a group-level centroid was first identified by running a canonical GLM analysis that compared the stimuli against their baseline (see Table 1) and then locating the peak of a statistical parametric map within the general areas associated with that CMC component (Fig 1). Because all tasks show stronger activation in the left hemisphere than in the right, all the group-level centroids were in the left hemisphere.

To account for individual-level variability in functional neuroanatomy, the group-level coordinates were then used as the starting point to search in 3D space for the closest activation peak within each individual statistical parameter map. Figure 7 illustrates the distribution of the individual coordinates of each region for each task, overlaid over a corresponding group-level statistical map of task-related activity (as in [29]). Each individual coordinate is represented by a point; the ∼200 points form a cloud that captures the spatial variability in the distribution of coordinates. Next, the individualized ROI coordinates were used as the center of a spherical ROI. All voxels within the sphere whose response was significant at a minimal threshold of *p* < .50 were included as part of the ROI. Figure

**Fig 7.**
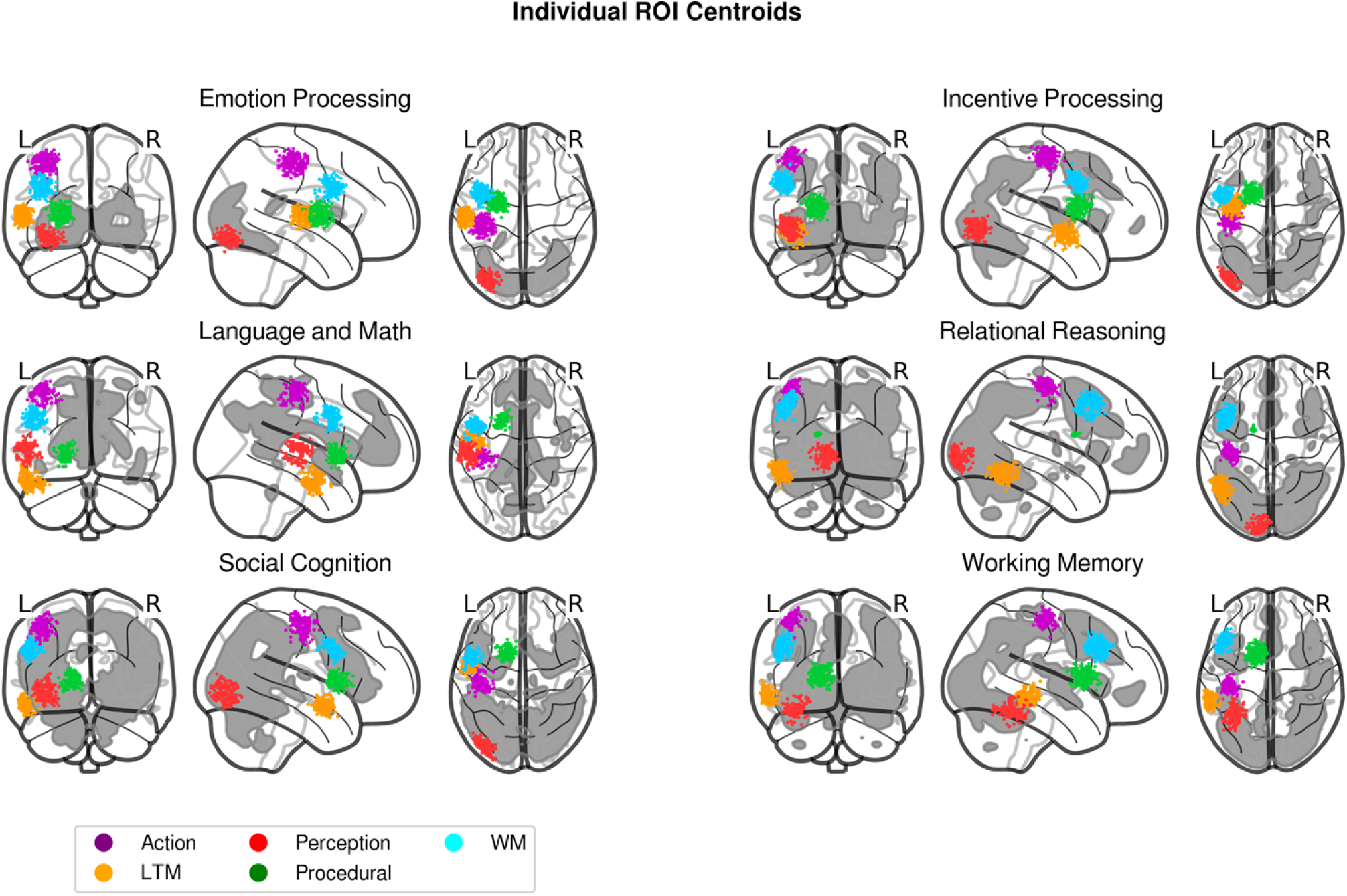
Location of ROI centroids across the six tasks of the Human Connectome Project. The colored dots, following the convention of Figure 1, represent the locations of the CMC components in every participant with variability in the location accounting for individual differences in functional anatomy. The grey background represents the group-level statistical parametric map of the main effect in each task thresholded at a statistical value of T > 5.

Finally, for each ROI of every participant in every task, a representative time course of neural activity was extracted as the first principal component of the time series of all of the voxels within the sphere.

#### Resting-state regions of interest

The method of ROI extraction outlined above relied on task-based activity to identify representative signals within large brain areas. However, this method depends on the assumption that any activity within a region is related to active completion of a task, which is not the case in resting state data. Instead, following [32], a meta-analysis method was used to more robustly identify potential regions associated with each of the Common Model components, and representative signals were extracted from these (Figure 8).

**Fig 8.**
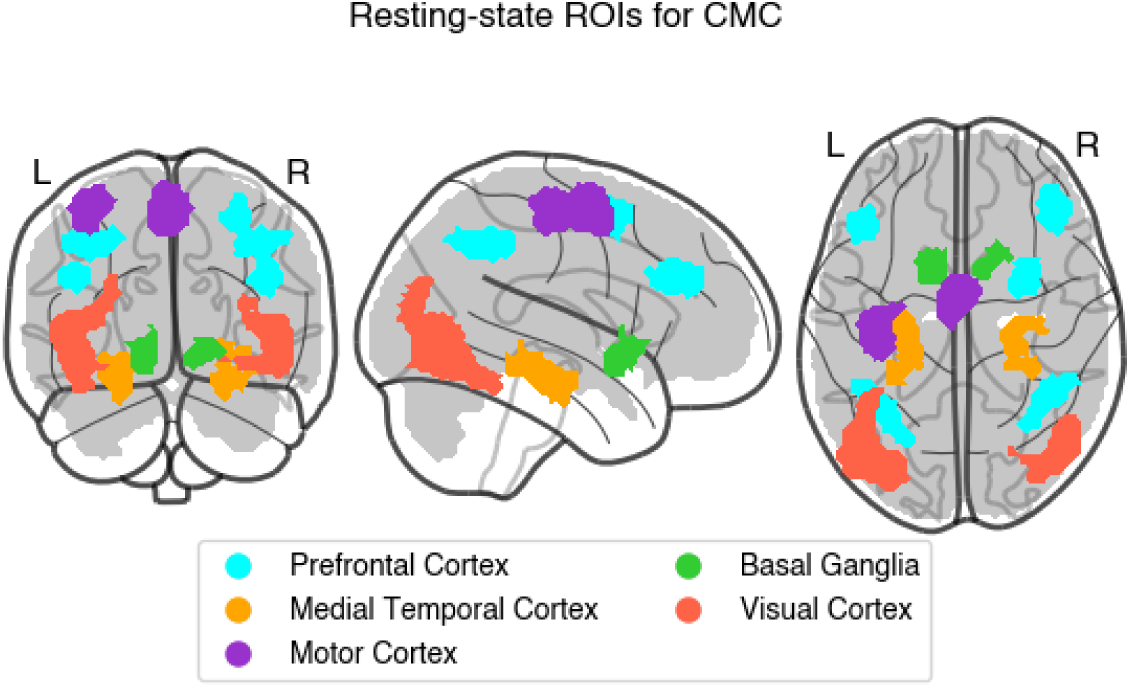
ROI regions for resting-state analysis. These regions were derived from a meta-analytic review of the literature [31] and were maintained for all participants. The grey background represents the group-level statistical parametric map of whether the variance captured by the oscillatory regressors (Fig. 6B; See [32]) was significantly greater than the variance of the residuals, thresholded at T > 5.

#### Model Fitting

Once the time-series for each ROI was extracted, different networks were created by connecting all of the individually-defined ROIs according to the specifications of each model (Figure 1). The predicted neural activity for each model was then calculated using Equation 1, and the predicted time course of the BOLD signal was then generated by applying a biologically-plausible model of neurovascular coupling to the simulated neural activity of each region. All of the model parameters were estimated through an expectation-maximization procedure [24] to reduce the difference between the predicted and observed time course of the BOLD signal in each ROI.

## Author Contributions

Conceptualization: Catherine Sibert, Andrea Stocco

Formal analysis: Catherine Sibert, Holly S. Hake, Andrea Stocco

Funding acquisition: Andrea Stocco

Investigation: Catherine Sibert, Holly S. Hake, Andrea Stocco

Methodology: Catherine Sibert, Holly S. Hake, Andrea Stocco

Project administration: Catherine Sibert, Andrea Stocco

Software: Andrea Stocco

Supervision: Andrea Stocco

Validation: Andrea Stocco

Visualization: Catherine Sibert, Holly S. Hake, Andrea Stocco

Writing – original draft: Catherine Sibert, Holly S. Hake, Andrea Stocco

Writing – review & editing: Catherine Sibert, Holly S. Hake, Andrea Stocco

## Data Availability Statement

All of the software code, statistical analysis code, DCM analysis files and data, and figures for this paper are available at https://osf.io/2gjwv/.

